# The variability of evolvability: properties of dynamic fitness landscapes determine how phenotypic variability evolves

**DOI:** 10.1101/2025.07.02.662714

**Authors:** Csenge Petak, Lapo Frati, Renske M. A. Vroomans, Melissa H. Pespeni, Nick Cheney

## Abstract

The magnitude and shape of phenotypic variation depends on properties of the genotype-to-phenotype (GP) map, which itself can evolve over time. The evolution of GP maps is particularly interesting in variable environments, as GP maps can evolve to bias variation in the direction of past selection, increasing the evolvability of the population over time. However, the degree and manner in which environmental variation shapes GP maps and influences evolutionary dynamics may depend on properties of the fitness landscape. To explore how evolutionary dynamics are affected by variable environments across a wide range of different pairs of fitness landscapes, we evolved GP maps to produce spatial-temporal gene expression patterns that matched two-dimensional patterns generated by different elementary cellular automata (CA) rules. We found remarkable variation in how populations evolved in variable environments. In some cases, changing the environment helped populations find higher fitness peaks; in others, it hindered them. The evolution of evolvability also depended on the fitness landscape pair. In some experiments, the ability to generate adaptive phenotypic variation upon environment change increased over time, while in some others, populations found shared areas between fitness landscapes. On the other hand, environmental variability consistently resulted in higher fitness landscape exploration, average fitness and mutational robustness compared to evolution in static environments, which we hypothesize are tightly connected. In conclusion, work presented here sheds light on important general consequences of environmental variability, while also demonstrating dependency on properties of fitness landscapes, which future research on the evolution of evolvability should consider.

**Significance statement:** The speed and direction of evolution depend on the availability of phenotypic variation. Genotype-to-phenotype maps can over time bias phenotypic variability to more readily produce alternative adaptive phenotypes in fluctuating environments. However, because properties of the fitness landscapes influence evolutionary dynamics, it remains unclear which previously observed dynamics reflect general effects of environmental variability and which are specific to the pair of landscapes used. We found that the height of the fitness peaks discovered, and how the populations became more evolvable, significantly differed across landscape pairs. In contrast, environmental variability consistently increased average fitness and mutational robustness. Thus, future research investigating the inherent consequences of frequent environmental change should be done on a range of dynamic landscapes.

## Introduction

Understanding adaptation to variable environments, how past evolution influences future adaptation, and how novel phenotypic variation comes to be are major open questions in evolutionary biology. Whether a particular population can adapt to a new environment or not depends on the ability of the biological system to produce new, useful phenotypic variation. The kind of phenotypic variation that is generated is not “random” (see [1] for an overview). In particular, the genotype-to-phenotype (GP) map influences the magnitude of variation along different dimensions of the phenotype in response to genetic and environmental perturbations. The developmental “constraints”, “biases” or “structure” [2, 3], influence what is easy to evolve and therefore what path evolution likely takes [4–6].

Importantly, the GP map can itself evolve. GP maps that produce more variation in the direction of selection will be favored; as the most adaptive phenotypes are selected, indirectly, the mechanism and mutational neighborhood that produced offspring with that phenotype is also selected [7, 8]. The evolution of GP maps is particularly striking in variable environments [9]. In a changing environment where there is some structure to the new pressures that appear, populations can evolve to become faster at generating adaptive phenotypic variation over time [4, 7, 10, 11]. This is because GP maps can evolve to bias phenotypic variation in order to produce phenotypes that are fit in alternative environments with fewer mutations, reducing the time needed to adapt to that environment.

The conditions and ways in which evolvability evolves in variable environments have been investigated both in experimental and theoretical work [12–17]. However, most studies consider only one scenario, one pair of alternative environments the populations need to navigate and get better at evolving in (e.g., [10, 18, 19], with a few notable exceptions such as [20, 21]). It remains unclear how environmental variability shapes evolutionary dynamics across different pairs fitness landscapes, and to what extent patterns in phenotypic variability, robustness, and evolvability are general or specific to particular landscapes. We aim to address this gap.

### Cellular automata patterns

To this end, we developed an agent-based model to simulate populations evolving in static and variable environments where each individual possesses a Gene Regulatory Network (GRN) with continuous weights that govern gene expression in a row of cells. The fitness of an individual depends on the expression level of a specific gene (here, *target gene*) in each cell throughout developmental time. To assess fitness, gene expression patterns across cells and through development were compared to target expression levels generated by different elementary cellular automata rules.

Cellular automata (CA) are often used as models of emergence of complex patterns from simple, local rules [22]. Elementary CA (ECA) update each element in a sequence of binary values using a deterministic rule. The CA rule determines the next value of each cell using that cell’s and its immediate neighbors’ state. A CA rule can be described using just 2^3^ = 8 bits, one for each possible configuration of 3 consecutive binary cells (see Fig. 1 as an example), for a total of 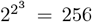 different possible ECA rules.

**Figure 1.**
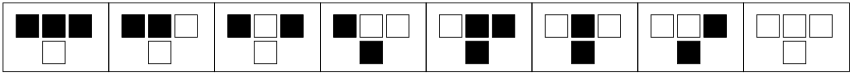
Top row: all possible input states for ECAs, 1 or 0 state of the cell and left and right neighbors. Bottom row: state of the cell after the update for rule 30. ECA rules are commonly referred to with a value between 0 and 255, here is an example of how to convert the rule above to its base 10 value: □□□■■■■□ = 000011110_2_ = 30_10_

Starting from a specific initial condition (i.e., sequence of binary values that is the input to the CA; t=0) a two-dimensional pattern can be generated by repeatedly applying a given CA rule a fixed number of times to each cell (columns) while keeping track of each new state (rows, Fig. 2 example patterns). The initial condition used to generate the specific target pattern for fitness calculation was also used as the starting point of the individual’s development. Thus, populations needed to evolve GRNs that created the target pattern given that the *target gene’s* expression level was set to the correct initial condition in each cell. The fitness function included the expression level of the *target gene* throughout development. This way, we were evolving a developmental process rather than a specific gene expression state to arrive to. Furthermore, this expanded the number and range of difficulty of possible targets to use for the evolutionary algorithm.

**Figure 2.**
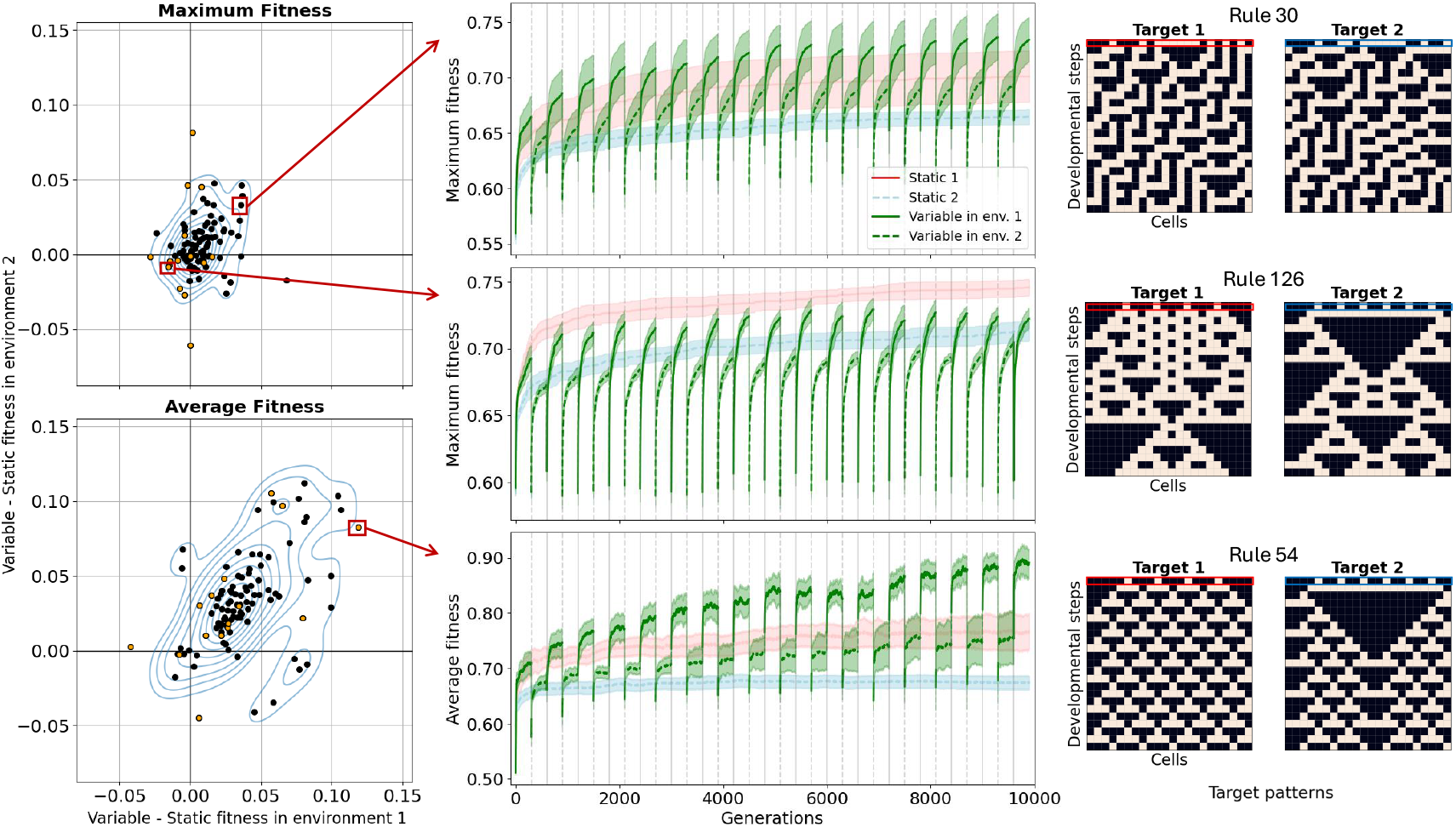
Left: Scatter plots showing the difference in highest average (bottom) and maximum (top) fitness between populations evolving in variable and static environments. X-axis compares fitness in environment 1 to evolution only in environment 1, y-axis compares fitness in environment 2 to evolution only in environment 2. Each of the 105 pairs of fitness landscapes is shown, with orange points highlighting the fitness landscape pairs chosen for more detailed analyses. There is large variation among the experiments and while the effect on maximum fitness is centered around 0, environmental variability has a positive effect on average fitness. Middle: Line plots highlighted by red arrows show maximum and average fitness over time for static and variable runs as examples. Top line plot shows an example fitness landscape pair where the variable runs had higher maximum fitness than runs in either of the static environments - which is why it is in the upper right quadrant of the scatter plot. Right: Pair of target patterns used for the experiments in the corresponding line plots. Targets within each pair were generated using the same CA rule with different initial conditions (highlighted by red and blue boxes).

In each experiment, environmental change was modeled as the periodic switching between two alternative target patterns that determine the fitness of individuals’ phenotypes. Two target patterns within a pair were always generated using the same CA rule but starting from different initial conditions (examples in Fig. 2). In total, 105 pairs of target patterns were generated by making deterministic pairs of 14 initial conditions and applying 15 different rules to each of them. Since the starting point of development changed with the target pattern and the CA rule was kept the same within an experiment, GRNs were evolving in alternative fitness landscapes that had both independent and shared areas. This is because in each pair, populations could get better at generating only the current pattern, or, by approximating the CA rule, both of the patterns simultaneously. Thus, different target patterns within a pair can be thought of as different instances of applying a general mechanism, and populations could get better at generating only one or both of the instances. This is analogous to applying the same developmental mechanisms in different contexts to get new phenotypes. We conducted experiments where the CA rule changed instead of the initial condition between the alternative targets - the main conclusions of this study apply to that setting as well.

Using this approach, we created a suite of pairs of fitness landscapes that had within and between landscape pair variation in the complexity of the target patterns resulting in varying levels of ruggedness and difficulty. We found that the effect of environmental variability on maximum fitness depended on the fitness landscape pair - sometimes it helped, while in other cases it prevented the populations from reaching higher fitness peaks. We hypothesize that this is because while in some settings environmental variability helped populations escape narrow local optima, in others, evolution in one environment “deceived” the population into worse areas of the fitness landscape with respect to the other environment. Additionally, we found significant differences in the ways populations became more evolvable across the different pairs. In some pairs populations evolved to be good in both environments, while in others they evolved to have GP maps such that a few mutations changed the phenotype from one target to the other. Thus, we found the biasing of phenotypic variation reported in previous literature only in certain landscape pairs. Lastly, we found that mutational robustness, average fitness and landscape exploration consistently increased across variable experiments compared to static ones. We tie these general effects together by suggesting that environmental variability promotes the discovery of wider fitness peaks across fitness landscapes.

## Results

### Evolution in static environments

We first explored the evolutionary dynamics of populations evolving in the 210 (15 rules with 14 initial conditions) static fitness landscapes. All target patterns can be found in Supp. Fig. 7. The fitness distribution of randomly initialized populations (t=0) varied substantially in mean, modality, and spread between the different fitness landscapes (Supp. Fig. 1.B). There was also variation in the maximum (range: 0.63 - 1) and average fitness (range: 0.54 - 0.99) after 10,000 generations. The maximum and average fitness in all runs plateaued by this point. The average and maximum fitness of both random and evolved GRNs depended more on the CA rule than the initial condition (Supp. Fig. 2). Overall, the selective environment (target pattern) strongly determined the ability of populations to adapt.

### Evolution in variable environments

The remainder of the Results section is organized into two parts. First, we present how maximum and average fitness change over time in variable environments and examine what factors explain variation in these metrics across different fitness landscape pairs. Second, we analyze within-population variation in fitness, phenotypes, and genotypes to investigate the dynamics and speed of fitness recovery after a change in the environment.

### Environmental variability has variable effects on maximum and average fitness

We evaluated the effect of environmental variability on the highest maximum and average fitness the population could reach in each of the 105 experimental set ups. This gave us an insight into the height of the fitness peaks the evolutionary algorithms could find, as well as the highest the average fitness of the population could get throughout the generations without the confounding factor of reduced fitness after an environmental change in variable experiments. To do this, for each pair of fitness landscapes, we first averaged across replicate experiments, then took the difference between the maximum values of variable and static runs throughout the generations for both maximum and average fitness (Fig. 2). The top-right quadrant in the scatter plots contains experiments with fitness landscape pairs for which the variable treatment resulted in a higher fitness compared to populations evolved in either static environments. Similarly, the bottom-left quadrant contains experiments for which populations evolving in either of the static environments outperformed the variable treatment. The remaining two quadrants contain experiments for which the variable treatment had a higher fitness compared to the population in one static environment but not the other.

We found that the effect of environmental variability differed between different pairs of fitness landscapes, since there were pairs of fitness landscapes in each of the four quadrants. To exclude the possibility that this variation was due to noise between replicate runs in the same experiments, we calculated statistical significance for each variable versus static experiment comparison within each pair of fitness landscapes and adjusted the p-values for multiple tests. The general pattern in Fig. 2 remained the same after dropping experiments with nonsignificant differences between variable and static runs.

### No general trend in maximum fitness

Environmental variability alone did not have a clear effect on the maximum fitness that the population could reach (Kruskal-Wallis H-test *p* = 0.055 with respect to environment 1, *p* = 0.175 with respect to environment 2.) However, the results clustered by CA rule (*p* = 0.0184 and *p* = 0.00473) and when accounting for that variation by using a two-way ART ANOVA, we found that environmental variability had an effect with respect to both environments (*p* < 0.001 for both). This suggests that varying the environment had an effect on maximum fitness, but this effect depended on the CA rule. This can be clearly seen from the upper scatter plot and example line plots in Fig. 2. For example, in the top line plot variability helped, in the middle it hurt the maximum fitness consistently across replicates. Thus, in some fitness landscape pairs environmental variability helped the population reach higher fitness, while in others, it hurt the population in terms of maximum fitness reached.

### Variability increases average fitness

Environmental variability had a clear, positive effect on the average fitness of the population (see Fig. 2 bottom scatter plot upper right bias in density). The CA rule alone still had an effect (*p* < 0.001 both), but so did variability. Combining all experiments together and calculating the effect of varying the environment alone showed an increase in average fitness in variable environments with respect to both alternative environments (*p* < 0.001 for both). Thus, while the effect of environmental variability on average fitness still depended on the CA rule, it had an overall positive effect across the fitness landscape pairs.

The above results generally held for experiments where instead of the initial conditions, the CA rules changed within a pair of target patterns. In these experiments, since the initial condition remained the same upon environment switch, the individuals were not given any signal about the change, and the mechanism for generating the target patterns couldn’t be reused across the environments. Again, we observed large variation, with experiments in all four quarters (Supp. Fig. 4). Additionally, average fitness was higher in variable environments (*p* < 0.001), while maximum fitness generally slightly decreased (*p* < 0.001). This means that the main conclusion that the effect of environmental variability is fitness landscape pair dependent and that environmental variability helps average fitness holds even when the populations are blind to environmental change.

As opposed to CA rule, initial condition had no significant effect on the maximum (*p* = 0.173) or average (*p* = 0.902) fitness the populations in variable environments reached compared to static ones. However, we found one pair of initial conditions (where the ON cells were evenly spaced, see Methods) where different pairs of fitness landscapes generated by different CA rules resulted in large variation in maximum and average fitness (colored orange on Fig. 2). We therefore focused our more detailed analyses on these 15 experiments.

### Features of target patterns did not explain variation in fitness effects

To look for possible explanations for the variation we observed in the effect of environmental variability, we first looked at features of the target patterns. We found no obvious feature of the target patterns that correlated with the effect environmental variability had on the maximum fitness the population could reach, with no effect of the number of cells ON in the targets, the percentage overlap between the targets, or the “difficulty” of the targets (defined as the average fitness that the static evolutionary runs could reach). On the other hand, we did find a negative correlation between the difficulty of the targets and the effect of environmental variability on average fitness (*r*^2^ = 0.097, *p* = 0.0012). So, the lower the average fitness of populations evolved in static environments, the higher the fitness gain from the variability. However, this correlation was weak and only explained a low percentage of the variation in the effect of variability. Thus, features of the target patterns don’t explain our conclusions about the variable effect of environmental variability.

### The benefit of random population displacement did not explain variation in fitness effects

It has been argued in previous literature that environmental variability could help populations reached higher fitness values because they escape bad local optima. Therefore, we then compared the effect of environmental variability to other ways of displacing populations from potential local optima, such as periodically adding random mutations or selecting in random directions. Although there was always an immediate large drop in fitness after applying the mutations (every 300 generations, same frequency as the one used for the variable experiments), in all but two of the 15 fitness landscapes, the maximum and average fitness of the population increased over time more than in the control static experiments. We also tested the effect of randomly moving the population in the fitness landscape by evolving the populations to match a randomly generated target pattern for 3 generations every 300 generations. Results from these experiments were very similar to the results obtained from randomly applying mutations. As such, compared to environmental variability, randomly moving the populations in the fitness landscape had a much more predictable and consistently positive effect. Furthermore, we investigated whether the fitness landscape pairs that benefited from environmental variability were also the ones that benefited most from other ways of potential escaping of local optima. We found no correlation between the effect of variability and random disturbance, meaning that fitness landscapes that benefit most from increased noise in exploration are not necessarily the ones where environmental variability helped the most. Supp. Fig. 5 shows examples of cases where environmental variability can significantly increase or decrease fitness while random movement had a moderate positive effect. In sum, environmental variability has a more complex and fitness landscape dependent effect then simply escaping local optima.

### Fitness variation at the end of evolution in the first environment did explain variation in fitness effects

For our last attempt to explain why environmental variability helped in some cases while it hurt in others, we investigated how well positioned the population was on the fitness landscape after evolution in one environment with respect to the environment to-be. To this end, we assessed the fitness effects of randomly mutating the highest fitness GRN immediately prior to an environmental change, relative to both the current and the new environment. We found large differences based on whether environmental variability helped or hurt fitness in the second environment. We found that while a positive correlation between fitness in the two alternative environments did not correlate with improved fitness in response to environmental variability (*r*^2^ = 0.0892, *p* = 0.279), a higher variation in fitness with respect to the fitness landscape to-be (after removing clear outlier rules 50 and 70, discussed later) was a strong predictor instead (*r*^2^ = 0.502, *p* = 0.0067). Focusing on the two fitness landscape pairs where environmental variability had the most pronounced effects on maximum fitness in the second environment, in the scenario where variability helped (rule 54), the population found itself on a “slope” in the new fitness landscape where a large proportion of the mutations resulted in higher fitness. On the contrary, in the scenario where variability hurt (rule 122), the population started evolution in the new fitness landscape on a wide, low fitness plateau as the vast majority of mutations had no effect on fitness (Supp.Fig. 6). Depending on the fitness landscapes, evolution in one environment could “deceive” the population to be in a region of the landscape from which evolution in the new environment was harder than from a randomly initialized population. This was the only predictor we found for the effect of environmental variability on fitness.

In the following, we present results pertaining to changes in fitness, phenotypes and genotypes over time across fitness landscape pairs to investigate the possible ways in which populations evolve in variable environments.

### The distribution of fitnesses evolve differently across fitness landscapes

To look for evidence of the evolution of evolvability we analyzed the fitness of individuals in the population over time and upon environment switch. As expected and shown in Fig. 2 and in Fig. 3, the typical dynamics of average and maximum fitness was an initial increase, then drop upon environment switch, followed by an period of increase again. In each season, defined as the period between environmental changes, the fitness increased above the levels reached in the previous season with the same environment, until reaching a plateau the population returned to after subsequent environment changes. At the same time, while the drop in fitness after environment switch increased in most experiments over time due to increased population fitness (Fig. 3.C), the fitness values the populations drop to also increased (Fig. 3.B). So, populations were “at a better starting point” each season start, especially compared to the first time they encountered the environment. Beyond this typical pattern, in some fitness landscape pairs the drop in fitness upon season change decreased over time or recovered in fewer generations, indicative of finding shared fitness peaks or areas where the two targets were fewer mutations away.

**Figure 3.**
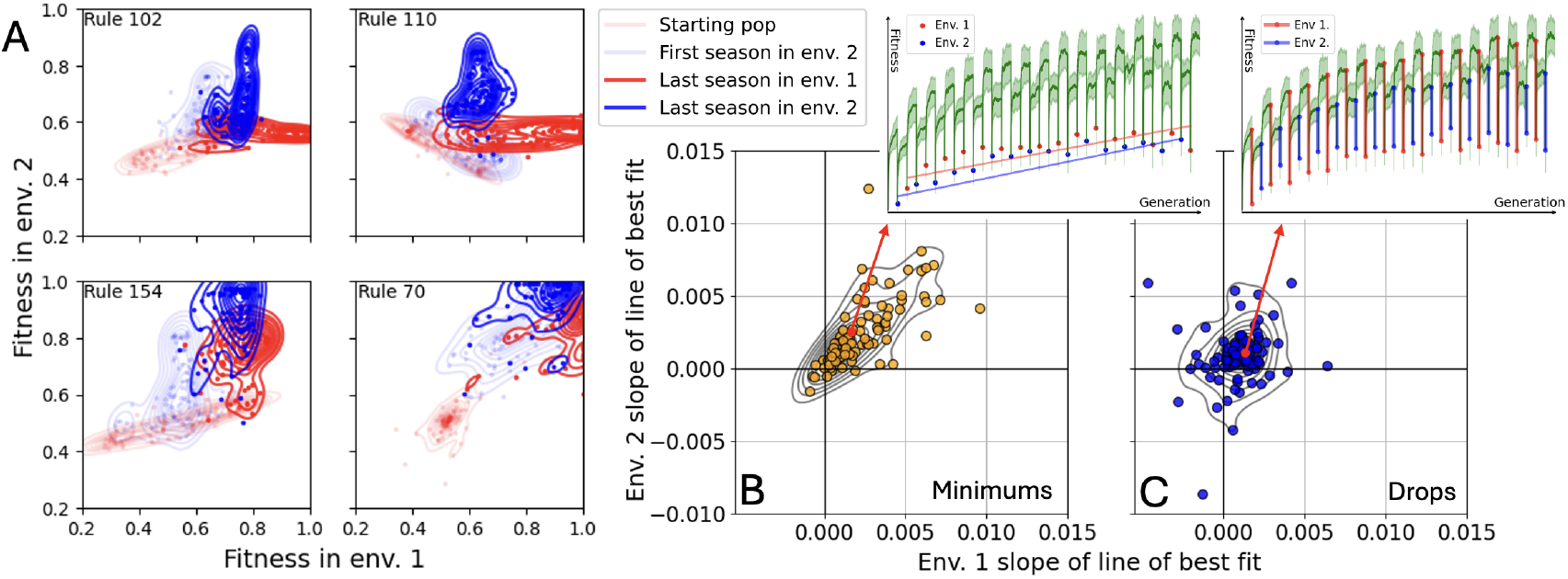
Population fitnesses over time. A) Fitness with respect to environment 1 (x-axis) and 2 (y-axis) at four timepoints during evolution (generation 0, 400, 9,500 and 9,800) for a random 10% of the population in four different experiments (different panels). Points represent specific individuals, lines show kernel density estimates. The evolution of fitness distributions varies substantially across different fitness landscape pairs. For the top two examples, individuals are only good in one environment at a time, while in the bottom two examples, the fitness distributions later on overlap, meaning that the populations find shared areas of the fitness landscapes. B), C) Each point is the average of 15 replicates for one of 105 experiments. X and y axes are the slope of the line of best fit of the average fitness in the population (yellow, B) or drop in fitness (blue, C) immediately after environment change over time. Same pattern holds for maximum fitness. Example experiment is shown in the insets. In both environments (env. 1 = red, env. 2 = blue) and in most experiments, the fitness value the populations drop down to increased (B), as well as the amount by which the fitness dropped (C). This means that the populations were at better and better starting points in the new environment, while the fitness dropped more overall as the populations reached higher and higher fitnesses by the end of a season. Still, in some experiments, fitness dropped less over time or recovered faster, suggesting the evolution of different forms of evolvability.

To further investigate these dynamics, in each generation we kept track of each individual’s fitness with respect to both environments to determine whether improvement in the current environment also increased fitness in the unseen environment. Overall, we observed two types of behaviors. In some experiments, fitness between individuals varied only with respect to the current environment (Fig. 3.A top row, “L” shape). In contrast, in other experiments, fitness variation aligned more over time, such that higher fitness individuals in one environment were also higher fitness in the other one (Fig. 3.A bottom row, diagonal and overlapping distributions). Thus, while in some fitness landscape pairs populations were constantly evolving to track the changing environment switching between peaks that are only good in the current environment, in others, populations were finding shared areas of the two fitness landscapes and could get better at both targets.

### Phenotypic variation is biased and reduced in response to environmental variability

The magnitude of phenotypic variation in the population and among offspring of the same parent depended largely on the fitness landscape, but in general, decreased through time. Moreover, across all fitness landscape pairs at the end of the experiments, GRNs from variable runs were more robust to random mutations than GRNs from static runs (Fig. 4.A). While there was variation among the 15 fitness landscape pairs, we found that the phenotype and fitness of mutated clones were more similar to the parent from variable environments than from clones of parents from either of the static environments (*p* < 0.001). Mutational robustness measured this way correlated with the amount of phenotypic variation measured in the population. Thus, mutational robustness evolved more in variable environments, even though robustness itself was never under direct selection in our model. This consistent pattern across fitness landscape pairs suggests that there are general consequences of evolving in variable environments.

Despite an increased robustness, we did find that the remaining phenotypic variation among offspring of the same parent was biased towards past selective environments. Components of the phenotype where the alternative target patterns overlapped varied less than components of the phenotype where the target patterns differed, see Fig. 4.B. For two of the fitness landscape pairs (54 and 30), the percentage of offspring with higher fitness than the parent at environment switch increased over time (Fig. 4.C), meaning that the populations were getting better at generating adaptive phenotypic variation - the evolvability increased. These two experiments were also the ones with the strongest bias in phenotypic variation towards the alternative target patterns. In these cases specifically, while shared areas between the fitness landscapes were not found, the populations were able to switch between alternative peaks increasingly fast.

**Figure 4.**
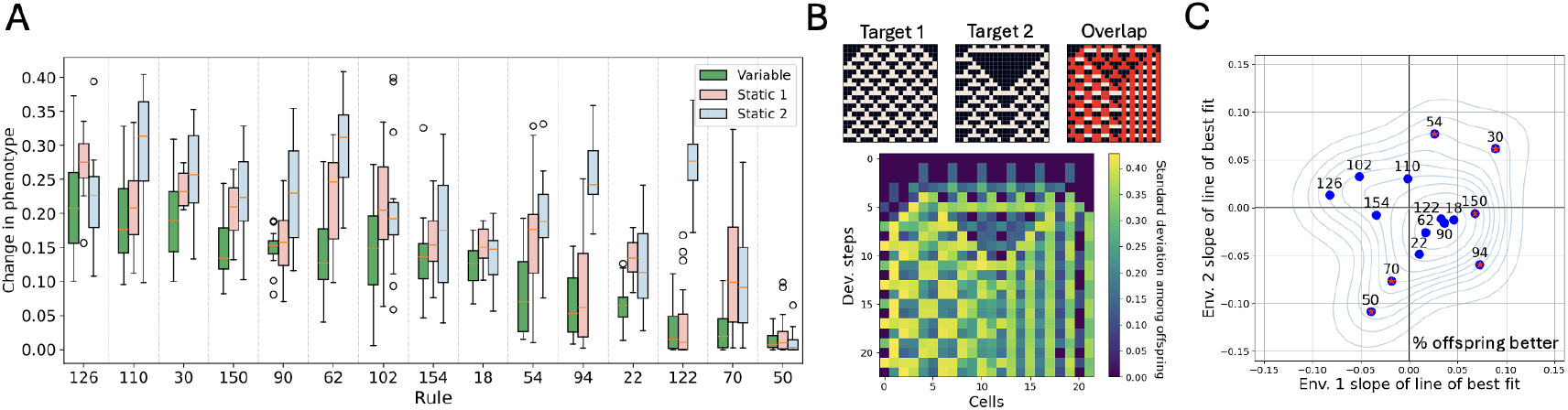
Phenotypic variability of populations evolving in variable environments. A) Average change in phenotype (where 0 is no change and 1 is exact opposite) as a result of random mutations of evolved individuals. Across the different experiments, individuals from variable experiments were more robust to mutations. B) Example of an individual with a GRN where the phenotype of mutated clones varies more in cells/ developmental steps where the target patterns differ. Top row shows target patterns and their overlap, bottom shows the standard deviation for each cell at each developmental step of phenotypes of mutated clones of a single individual. The evolution of this bias in phenotypic variability varied strongly with fitness landscape pair. C) Each point is the average of 15 replicates for each of the 15 experiments. X and y axes are the slopes of the lines of best fit of the percentage of offspring with higher fitness than the parent after each environmental change over time. For experiments 54 and 30 a larger percentage of offspring have higher fitness than the parent over time, hence them being in the upper right quadrant. In these two landscape pairs populations evolved to be fit in only one environment at a time, but they were faster at switching upon season change. Red stars indicate statistical significance.

On the contrary, in other experiments, shared areas were discovered as GRNs evolved to produce the new target pattern given the new initial condition and thus the same genotype could be high fitness in both environments. Additionally, some of these GRNs could generate the correct phenotype given previously unseen initial conditions, examples of this can be seen in Supp. Fig. 3. This happened in fitness landscape pairs 50 and 70 consistently across replicates. In these experiments, the drop in fitness decreased over time, phenotypic variation was not biased in a meaningful way, and the percentage of offspring with higher fitness than the parent decreased over time (Fig. 4.C). Populations evolving in these experiments followed a different path towards being adaptive in a variable environment and didn’t need new mutations to switch between alternative targets.

### Populations explore the landscape more in variable environments

Finally, we investigated how much genotypes changed during evolution; both in terms of how much the populations explored the landscapes and whether populations in variable environments ever got closer in genotype space to where they were before when they were in the same environment. We found that in all cases, populations in variable environments explored the landscape significantly more. GRNs at the end of the experiments were more different from the beginning (Fig. 5.C) and between replicate experiments (Fig. 5.B) in variable environments, as the genotypes changed through time much more than in static environments (Fig. 5.A). This means that even in cases where environmental variability hurt the maximum fitness the population could reach, as opposed to populations evolving in static environments, the populations were not stuck in a narrow local optima but instead kept changing over time. Interestingly, the genetic variation within a single population was similar in variable and static experiments.

**Figure 5.**
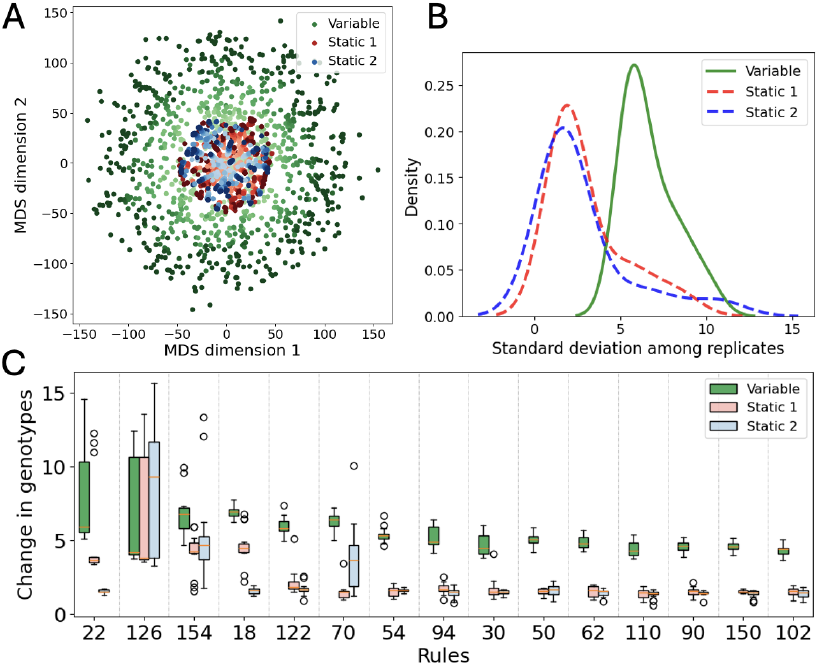
Genotypes (weights of the GRNs) changed more over time in variable environments. A) Multidimensional scaling of evolved GRNs show that populations explore the landscape more when the environment varies. Each point is the highest fitness GRN in the population at 33 equally spaced generations for 15 replicates for each treatment group in a representative experiment (rule 18). Darker colors indicate later generations. B) Standard deviation of evolved GRN weights among replicates is higher across different experiments in the variable treatment group (*p <* 0.001). C) The average difference between GRN weights at the beginning and end of an experiment in consistently higher in variable runs (*p <* 0.0001).

Additionally, we found that even though the environment populations experienced kept switching between just two alternatives, the evolved genotypes never got closer to where they were before. Both the distance from the first generation and the previous generation monotonically increased over time. Thus, they were not oscillating between the same two areas in genotype space as one might imagine in a low dimensional space, although the distance in genotype space increased less over time with respect to the last season with the same environment than the last season with the alternative environment. In summary, populations in variable environments continuously explored the high dimensional fitness landscapes across all experiments, which could tie into the other general effects of environmental variability we found regarding increased mutational robustness and average fitness.

## Discussion

In this study, we analyzed evolutionary dynamics across different pairs of fitness landscapes to investigate general and specific effects of environmental variability. Overall, we found a lot of variation across different fitness landscape pairs. The effect of environmental variability on maximum fitness varied significantly, as did the “strategy” to deal with variability. In some settings, populations found shared areas of the fitness landscapes where increasing fitness in one environment also increased fitness in the other environment. In contrast, we found that other populations improved at only one target pattern at a time but were getting faster at switching between phenotypes after the environment changed. Despite these differences, we also found strong general trends - populations in variable environments had consistently higher exploration of the fitness landscape, greater robustness to mutations, and higher average fitness compared to populations in static environments.

### Does environmental variability hurt or help fitness

Previous research on adaptation to environmental variability has yielded contrasting results, depending on the complexity of the specific problem space of investigation. In simple cases, the model assumes that evolution in the control static environments already finds the solution in a reasonable amount of generations (or computational time) and that varying the environment means that the populations have to continually re-adapt and find those peaks again and again, preferably in fewer and fewer generations as evolvability evolves [4, 10, 11, 20, 23]. Certain fitness landscape pairs examined in our experiments resembled this scenario. When the target patterns were simple (e.g., checker boards or stripes), global optima were found in both the static and variable runs. In these types of experiments, environmental variability has a negative effect if the environment changes faster than the speed at which populations can readapt and the population “turns around” before it can reach the same high fitness. In contrast, when exploring evolutionary dynamics in more complex environments, previous work found a benefit to varying the environment periodically. It is often argued that the dynamic changing of the fitness landscape can help populations escape local optima and explore the fitness landscape better [20, 21], resulting in an increased fitness [24, 25], or in a substantial speed up in terms of the number of generations needed to reach the same high fitness [20, 26].

Previous works paint a picture where variability hurts simple setting but helps complex ones. However, our experiments show that even in complex scenarios environmental variability can hurt by getting populations stuck on lower local optima (indicated by faster plateauing and worse fitness distribution of mutations of evolved genotypes). We hypothesize that in these complex cases where variability hurt, it was because evolution in one environment “deceived” the population with respect to evolution in the other environment. The shape of the fitness landscape is known to be important in determining a population’s evolutionary dynamics over time. If the landscape is rugged, populations can get stuck in local optima more readily than in smoother landscapes [20, 27]. However, ruggedness is only one proxy for the difficulty of a landscape. For example, problems with hierarchical structures can be more or less deceptive, in the sense that mutational trajectories that increase fitness might guide the population to local optima that are further away from global optima than the starting population [28]. Here, evolving in another environment first lead the populations to areas of the fitness landscape that they would not have ended up on otherwise, and in these cases the populations had a “worse start” than populations randomly generated in static experiments (see Fig. 6c for visual explanation). This is opposite to the scenario where evolving in one environment helps evolution in the other, a case depicted visually in [20] Fig. 5 and in [25] Fig. 1.

**Figure 6.**
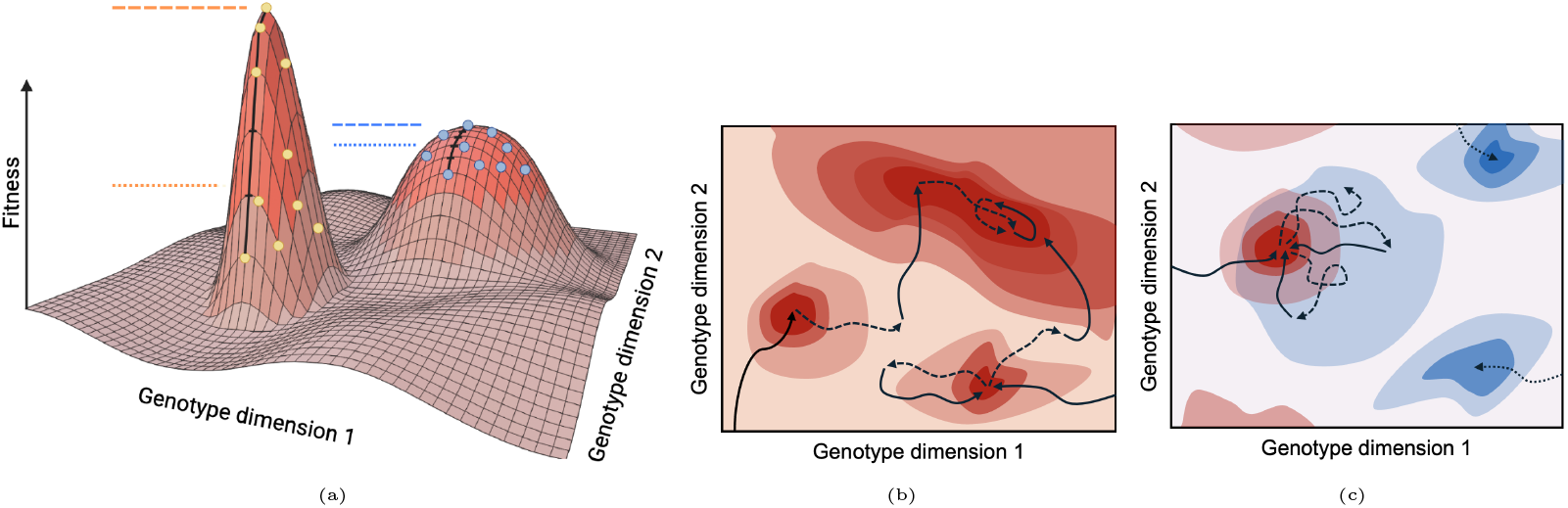
Schematic representation of fitness landscapes. **a)** Orange lines show maximum (dashed) and average (dotted) fitness of a population on a narrower peak (yellow circles), blue lines show the same for a wider peak (blue circles). A large difference between the maximum and average fitness of the population indicate a narrower local optima. **b)** Trajectory of a population in a periodically changing environment with respect to a given fitness landscape (solid arrow) and with respect to an unseen other fitness landscape (dashed arrow). Wider peaks are more likely to be found and stayed on. **c)** Hypothetical “trapping” of the population in variable environments. Evolution during the blue environment (dashed arrow) is stuck at lower fitness due to evolution during the red environment (solid arrow) and a better blue peak found by evolution in static environments (dotted lines) is thus never found in the variable environment.

### Does environmental variability result in increased evolvability

Evolvability can be defined and quantified in a range of different ways [13, 29–31]. Additionally, there are many ways in which a population can become more evolvable over time, for example, by increasing mutation rates or by evolving a way to bias the distribution of mutational effects toward beneficial phenotypes. Recently, it has been shown that both can evolve simultaneously in variable environments [18]. We investigated evolvability on the individual level by analyzing the magnitude and shape of phenotypic variability a specific genotype can produce given random mutations (Fig. 4), and on the population level by measuring the extent to which average and maximum fitness declines after a change in the environment (Fig. 3). Putting together the individual and population level investigations in this study, we found significant variation in whether and how evolvability evolved across different fitness landscapes. In some experiments, populations found areas of the fitness landscapes where evolution in one environment could increase fitness in both environments simultaneously. In these cases evolution in the alternative environments resulted in the closer approximation of the underlying mechanisms generating the target patterns (i.e., the CA rules), rather than a mechanism implementing a spurious correlation in only one of the target patterns (e.g., stripes). Often this has resulted in the ability to create the correct pattern even to previously unseen initial conditions.

However, in other experiments, the populations only evolved to be fit in the current environment. In a subset of these, populations consistently evolved to bias phenotypic variation generated by random mutations and the percentage of offspring with beneficial phenotypes after environment change increased over time. So, instead of finding shared areas, they found areas of the genotype space that were mapping to a high fitness in only one of the environments, but that were in each other’s mutational neighborhood such that upon a change in the environment the populations could find the new optima faster (see [18] Fig. 1.C for a visual depiction). Such evolution of evolvability has been found to evolve in many GRN studies before [10, 12, 16], however, here we found that whether this form of evolvability evolved was highly fitness landscape pair specific and in many experiments evolvability didn’t evolve according to our metrics.

The shape of the fitness landscape influences the likelihood of a specific phenotype to be found and maintained by evolution [32]. For example, the prevalence of a phenotype in genotype space has been found to be an important factor [33]. In a similar vein, we postulate that if areas of the genotypes space that are near boundaries of alternative high fitness phenotypes are abundant and easily accessible, evolvability in this form is more likely to evolve and stick around in the population. Thus, evolvability itself is more or less likely to evolve depending on the shape of the alternative landscapes [34].

### General effects of environmental variability

While we found no consistent effect of environmental variability on maximum fitness, variability generally helped the populations reach higher average fitness. In addition, we found an increased mutational robustness, calculated as the average difference between offspring and parent phenotypes, in experiments with environmental variability. The difference between the maximum and average fitness in the population is indicative of the width of the fitness peak they occupy, and consequently the mutational robustness of the individuals in the population. Two populations with the same maximum fitness can have different average fitness as more robust populations inhabiting wider fitness peaks display a smaller distance between maximum and average fitness [35] (see Fig. 6.a). Together, these results suggest that populations in variable environments were finding wider peaks.

The evolution of robustness is often observed in computational models of GRN evolution [36, 37], especially after an episode of fitness plateauing [2]. In this study, in most static experiments, populations climbed to the nearest narrow local optima and stayed there. In contrast, populations in variable environments explored the genotype space more. We hypothesize that this constant travel through genotype space kept populations away from narrow local optima. Finding a narrow peak in one environment meant that the population got stuck but only temporarily, since by the end of the following season the population likely moved off of the narrow peak. The wider the peak the population found the more likely it was that they were still on it, or at least near it, when the environment changed back (see Fig. 6.b). This could explain why populations in variable environments found wider optima and thus were more mutationally robust.

There is a rich literature on both the molecular mechanisms and the theoretical evolution of robustness [13, 35], and increased robustness has been observed in variable environments before [34]. However, our understanding is limited on how and why robustness evolves differently in variable and static environments, which future research should explore further [38]. We showed here that, in most cases, environmental variability also helped the populations reach higher average fitness, which could be the consequence of finding wider peaks while still being able to find not-so-bad height peaks as maximum fitness was less effected by environmental variability.

## Conclusion

We examined the effect of environmental variability on the evolution of evolvability and phenotypic variability across a range of fitness landscape pairs. We found that most measured metrics varied substantially and hypothesize that this variation arises from the interplay between different shapes and features of these complex fitness landscapes as populations transition between them. Furthermore, we found that average fitness, as well as mutational robustness and genotype-space exploration increased consistently in environmentally variable experiments. Taken together, these results indicate that the fitness landscape can heavily influence how much evolvability evolves. Future research on the evolution of evolvability and biased phenotypic variability should consider this effect.

## Methods

### The genotype

Following others [7, 39], we modeled the set of regulatory interactions between genes in the evolving gene regulatory networks as an adjacency matrix *W*, representing a weighted, directed graph that made up the genotype (*W*_full_). Every individual was made up of a strip of *C* cells with periodic boundary conditions (i.e., loop). Each cell within an individual shared the same GRN with *G* genes. Cells could communicate with one another; one specific gene was chosen whose expression influenced the left cell, and one other gene was chosen to influence the right cell (*communication genes*). Thus, *W*_int_ was made up of *GxG* weights to form a fully connected network where every gene regulates every other gene, and *W*_full_ was made up of *W*_int_ + 2*G* weights that connected the left and right neighbors’ communication genes to every gene in a cell. See Fig. 7A.

The expression levels of the genes were updated through the iterative multiplication of *W*_full_ by a vector of gene expression levels 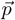 as follows:

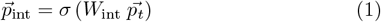

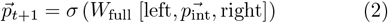

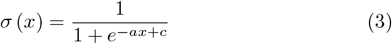

where σ is a sigmoid function (*a* = 10, *c* = 5) that constrains the value of 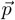 to be between 0 and 1. Step 1 uses the state of internal genes to create a temporary state that is then combined with information coming from neighboring cells in step 2.

### The phenotype

The phenotype of each individual was the gene expression level of a specific (noncommunication) gene over all the cells and steps of development to produce a two-dimensional grid of size [#cells ×# developmental steps]. In the following we will refer to the gene whose expression level contributes to the phenotype as *target gene*, and to genes that do not contribute to the phenotype as *internal genes*. Note that the leftmost and rightmost internal genes are used in communication and therefore are also called *communication genes* (see Fig. 7 for a visual explanation). At the beginning of development, each individual’s gene expression pattern was initialized to 0 for all genes except for the target gene in specific cells depending on the experiment (referred to as initial conditions). The fitness was calculated as the sum of absolute differences (Manhattan distance) between the phenotype and a target pattern generated by the repeated application of a CA rule, starting from the same initial condition as the developmental process. We could test how well the evolved GRNs internalized the CA rules by applying the same GRN to unseen initial conditions (e.g., in Supp. Fig. 3).

**Figure 7.**
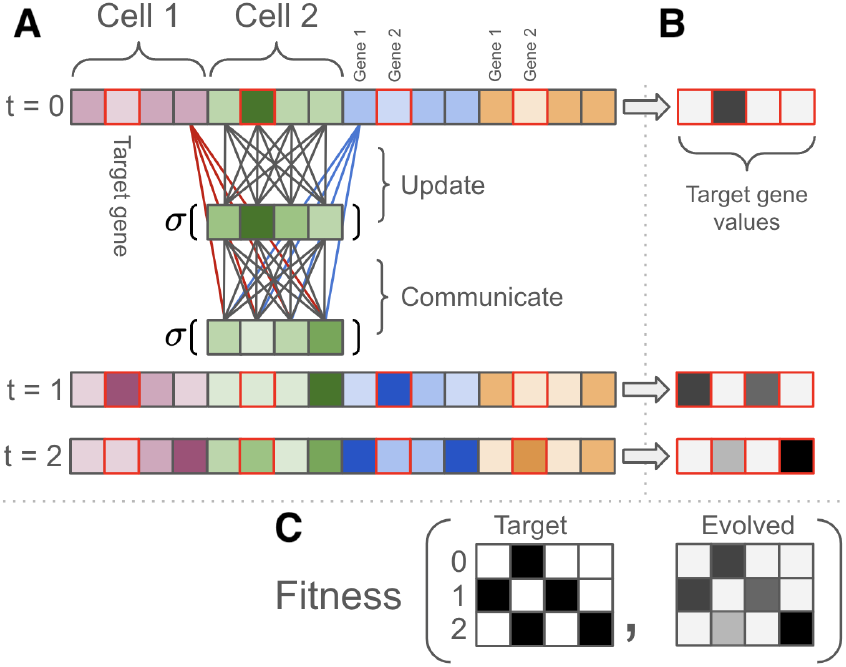
**A**: The same GRN (*W*_full_) updated the state of all cells within an individual. It contained *G* × *G* + 2*G* coefficients (4 × 4 + 8 in the example). To update the state of one cell the GRN received the gene values of genes in that cell (16 **grey** edges, *W*_int_) and two genes coming from that cell’s neighbors (4 **red**/4 **blue** edges). Grey weights are used both in the Update and Communicate steps, red and blue weights are used only in the Communicate step. **B**: The value of a specific target gene is extracted from all the cells (here the second gene, red border), the extracted values form a sequence of length equal to the number of cells. **C**: Target gene values across multiple development steps are combined and compared against the target patterns to compute the fitness of the individual.

### Evolutionary algorithm

A population of haploid, asexual individuals were generated with weights for *W*_full_ randomly drawn from a normal distribution (𝒩(0, 1)). The population was then evolved using a µ + λ Evolution Strategy with truncation selection. Individuals were sorted based on fitness and the top µ ones were selected to survive to the next generation, and generate (popsize/µ) ™ 1 offspring each to keep a constant population size. Offspring were mutated at *G* * *m* positions by adding a small random value drawn from a normal distribution (𝒩(0, 0.5)). Experiments were run for 10,000 generations, which was the point at which fitness values generally plateaued. Experiments run for 30,000 generations resulted in no new insights. See Supp. Table 1 for more detail on the parameters used in experiments.

In experiments with variable environments, the fitness function compared the phenotypes to two different target patterns, switched at a fixed rate periodically. These target patterns were generated using the same CA rule but with different initial conditions. We made a total of seven pairs of initial conditions, six of which were generated by randomly shuffling a list containing 11 ones and 11 zeros (*C* = 22 in every experiment) for each target in each pair, making sure that none of the twelve lists generated this way were the same under rotation. The seventh pair was created manually by arranging four versus six 1s equally spaced. This seventh pair was the one that was used for more detailed analysis - as described in the Results section. Many distinct patterns can be generated by changing either the rule or the initial condition, though in practice not all of the 2^#Cells^ times 256 patterns are unique as sometimes the same or rotationally equivalent patterns appear by applying different rules to the same initial condition or vice versa [40]. Thus, CA rules used in the experiments were chosen manually out of the 256 possible rules based on visual complexity of the produced patterns across the initial conditions; rules that produced duplicate or symmetrical pairs to the ones already created were discarded, along with rules that produced very simple patterns (e.g., all cells ON or all cells OFF). Supp. Fig. 7 shows all 105 pair of target patterns generated this way.

### Analysis of evolutionary dynamics

Maximum and average fitness was compared between variable and static runs for each fitness landscape pair using the Mann-Whitney U (scipy.stats, [41]) test followed by Benjamini/Hochberg calculation of False Discovery Rate (statsmodels, [42]) to account for multiple testing across fitness landscape pairs. Kruskal-Wallis H-test (scipy.stats) was used to calculate the significance of environmental variability alone, and a two-way ART ANOVA (statsmodels) was used to calculate the effect of combinations of variables (e.g., environmental variability together with CA rule on maximum fitness, average fitness, and robustness). Linear least-squares regression (scipy.stats) was used to find relationships between continuous variables.

Parameters for random mutation introduction experiments: Every 300 generations every individual was mutated at every connection in the GRN by a value randomly drawn from a normal distribution (𝒩(0, 1)). Parameters for random selection experiments: Every 300 generations a new random target pattern was generated by drawing a number at random from a normal distribution (𝒩(0, 1)) for each cell in the 2D pattern. This new target was used in the fitness function determining selection for 3 consecutive generations. Both of these experiments were repeated 5 times on 15 different fitness landscapes (second of each fitness landscape pair used in the experiments with environmental variability).

Exploring mutational neighborhood of an evolved GRN involved the generation of 2,500 clones that were mutated by adding noise to each connection drawn from a normal distribution. Phenotypic robustness was measured as the average absolute difference between the phenotype of the mutated clone and the original GRN across all the clones. Fitness robustness was measured as the average difference between fitnesses of mutated clones and the original GRN. Similarly, to assess phenotypic variability, 10,000 mutated clones were created to measure variation in phenotypes across cells and developmental steps. Finally, the multidimensional scaling of GRN weights to evaluate genotypes space exploration was performed using 2 components, scikit-learn [43].

## Author contributions

C.P. and L.F. conceived the study. C.P. performed the research, analyzed the data, and wrote the manuscript. L.F. contributed to model development, data interpretation, and writing. R.V. contributed to model development and manuscript editing. M.P. and N.C. provided conceptual input and editorial feedback.

## Author declaration

The authors declare no competing interest.

## Acknowledgments

This material is based upon work supported by the 2021-2022 University of Vermont Dr. Roberto Fabri Fialho Research Award to C.P. and the National Science Foundation (NSF) Research Traineeship, Quantitative and Evolutionary STEM Traineeship (QuEST; DGE-1735316), IOS-1943316 supporting C.P. and M.H.P., and OIA-2218063 supporting L.F. and N.C.. Computations were performed on the Vermont Advanced Computing Core supported in part by NSF Award No. 1827314.

## Supplementary Information

**Figure S1.**
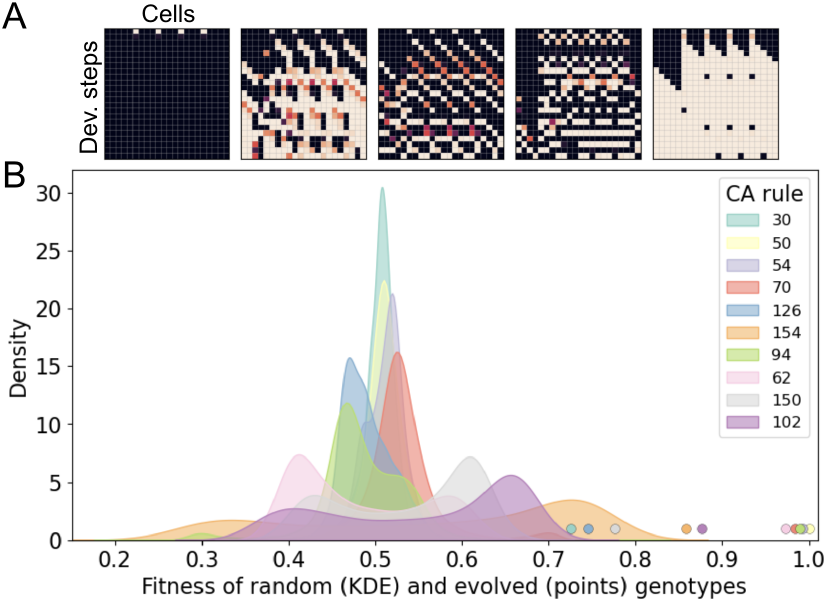
A) Phenotype of five randomly generated GRNs. B) Fitness distribution of randomly generated GRNs with respect to 10 different target patterns. Color coded points correspond to the maximum fitness of evolved GRNs in these fitness landscapes.

**Figure S2.**
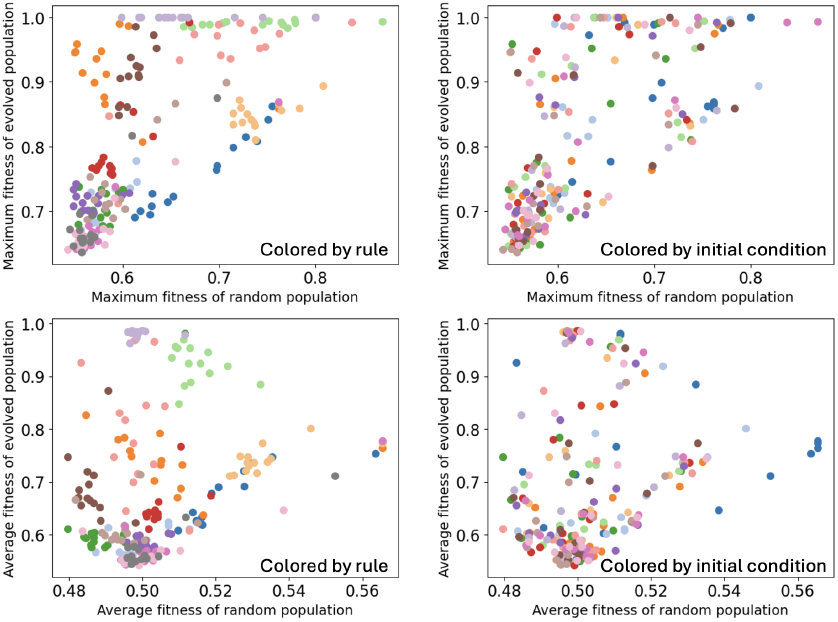
Maximum and average fitness of statically evolved and random populations averaged across 15 replicates for all 210 fitness landscape pairs. Left subfigures are colored by CA rule, right subfigures are colored by initial condition. Fitnesses cluster by CA rule more along both y and x axes. The standard deviation across experiments with different initial conditions is ∼3× higher than across different CA rules for evolved populations, and ∼1.5× higher for random populations, in terms of both maximum and average fitness.

**Figure S3.**
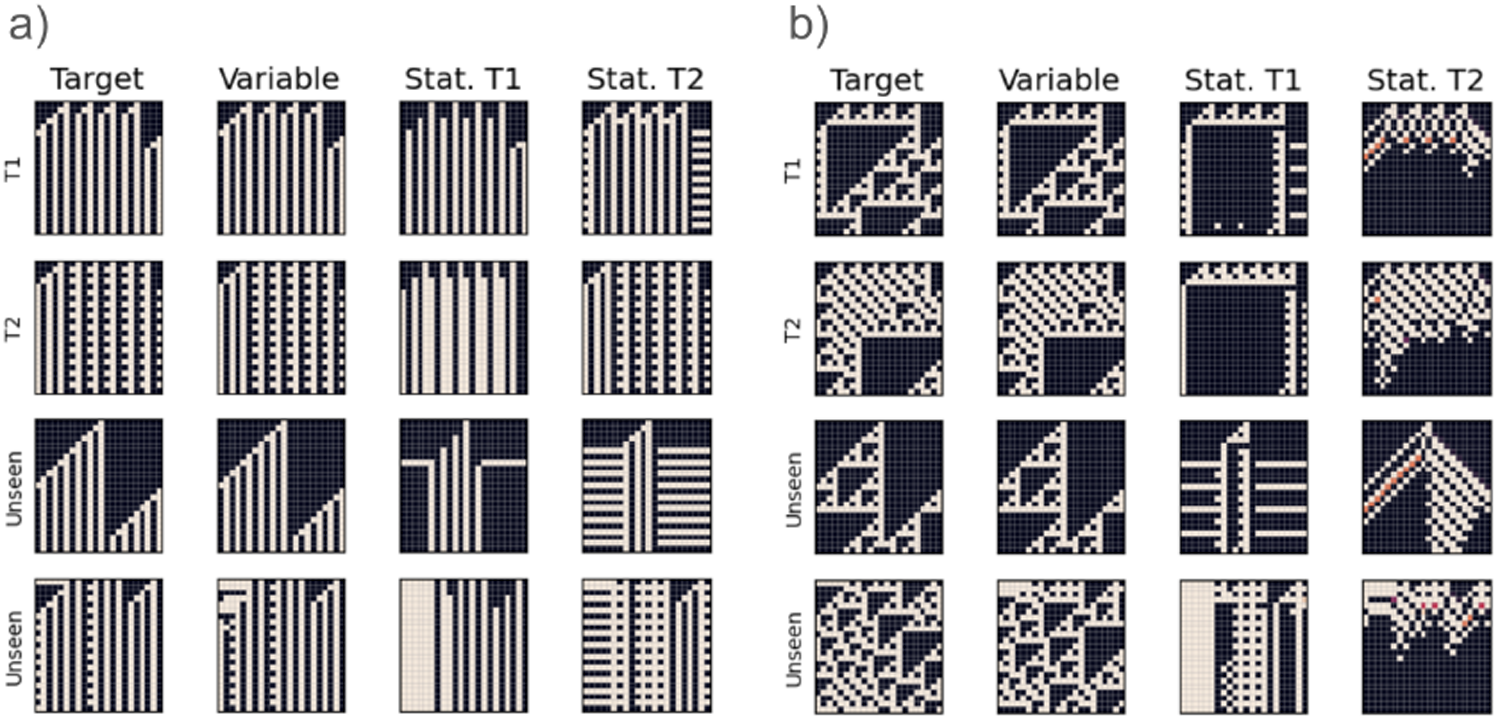
Example GRNs evolved to match patterns generated by rule 70 (a) or rule 102 (b). Rows: Output of the same GRN given different initial conditions. Columns left to right: Target pattern, GRN evolved in variable environment, GRN evolved in only the first environment, and GRN evolved in only the second environment. In these settings, GRNs evolved in variable environments were able to better match unseen patterns generated by the same CA rules.

**Figure S4.**
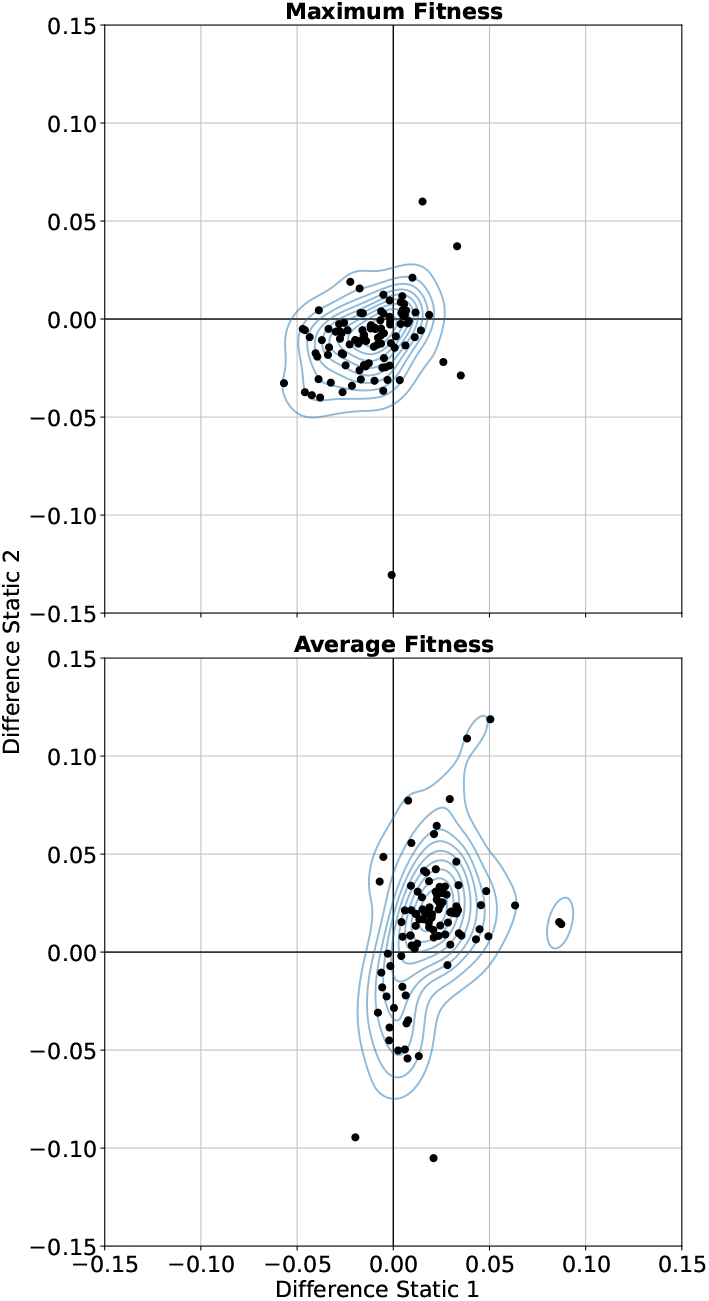
The effect of environmental variability on maximum and average fitness in cases where the alternative evolutionary target patterns were generated using the same initial conditions with different cellular automata rules. Thus, the alternative fitness landscapes didn’t share global optima. Everything else was kept the same such that results from Fig. 2 are comparable. Again, environmental variability has a positive effect on average fitness. Additionally, environmental variability has an overall negative effect on maximum fitness. Patterns hold when accounting for multiple testing.

**Figure S5.**
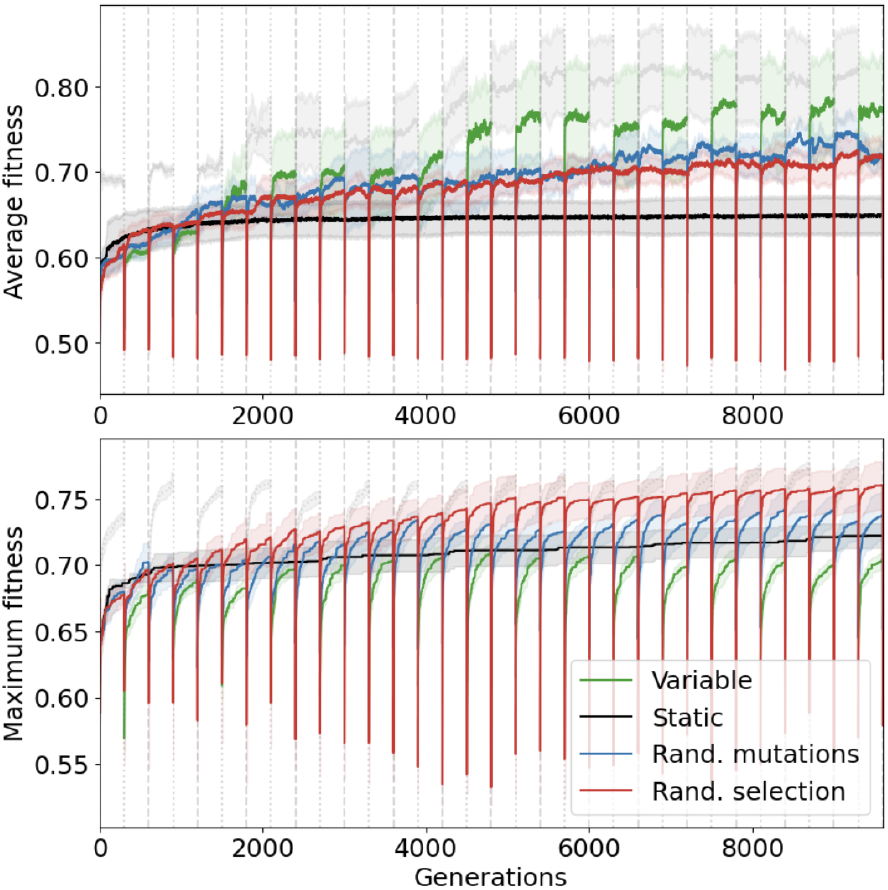
Average (example experiments with rule 102) and maximum (example experiments with rule 150) fitness in the population over time in variable and static environments, as well as in runs where the individuals were randomly mutated (blue lines) and in runs where the target pattern was randomized for three consecutive generations (red lines) every 300 generations. For runs with variable environments the line is grayed out for generations evolving to the alternative target for easier visual comparison. Both random mutation and random selection had a consistently positive effect over static experiments, while the effect of environmental variability was more variable.

**Figure S6.**
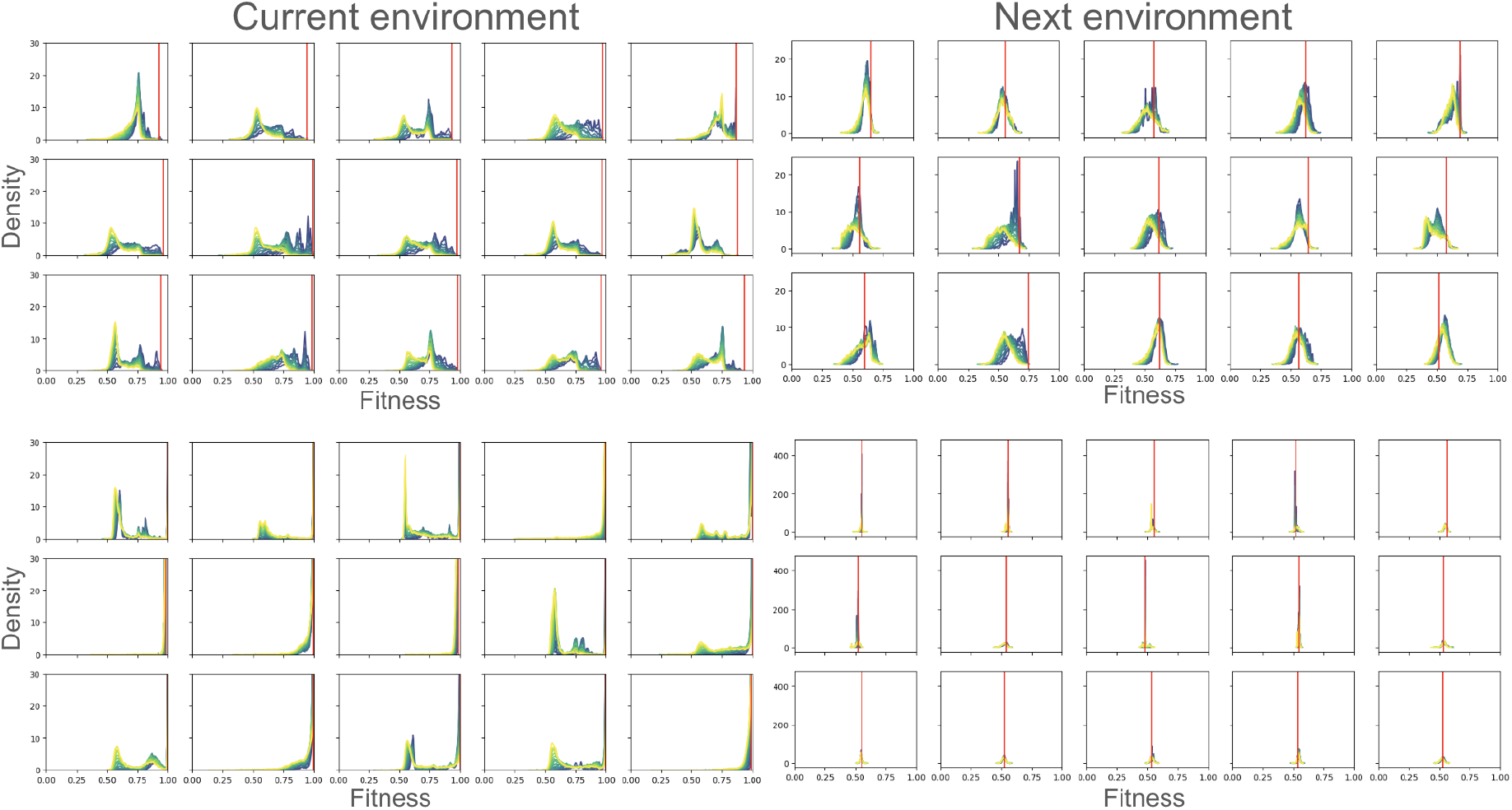
Distribution of fitness effects of mutations applied to the GRN with the highest fitness at the end of the first season (300 generations) across 15 replicate runs. Subfigures on the left show the fitness calculated with respect to the current, subfigures on the right with respect to the next environment. Top half shows replicate runs of experiments with rule 54, while the bottom half shows rule 122. Different colors show different mutation rates, ranging from 𝒩(0, 0.063) (blue) to 𝒩(0, 0.3) (yellow). Red lines show evolved GRNs’ fitness. While most mutations had a negative effect in both experiments with respect to the current environment, in runs evolved for rule 54 (top half) a lot of these same mutations were beneficial with respect to the environment in the next generation.

**Figure S7.**
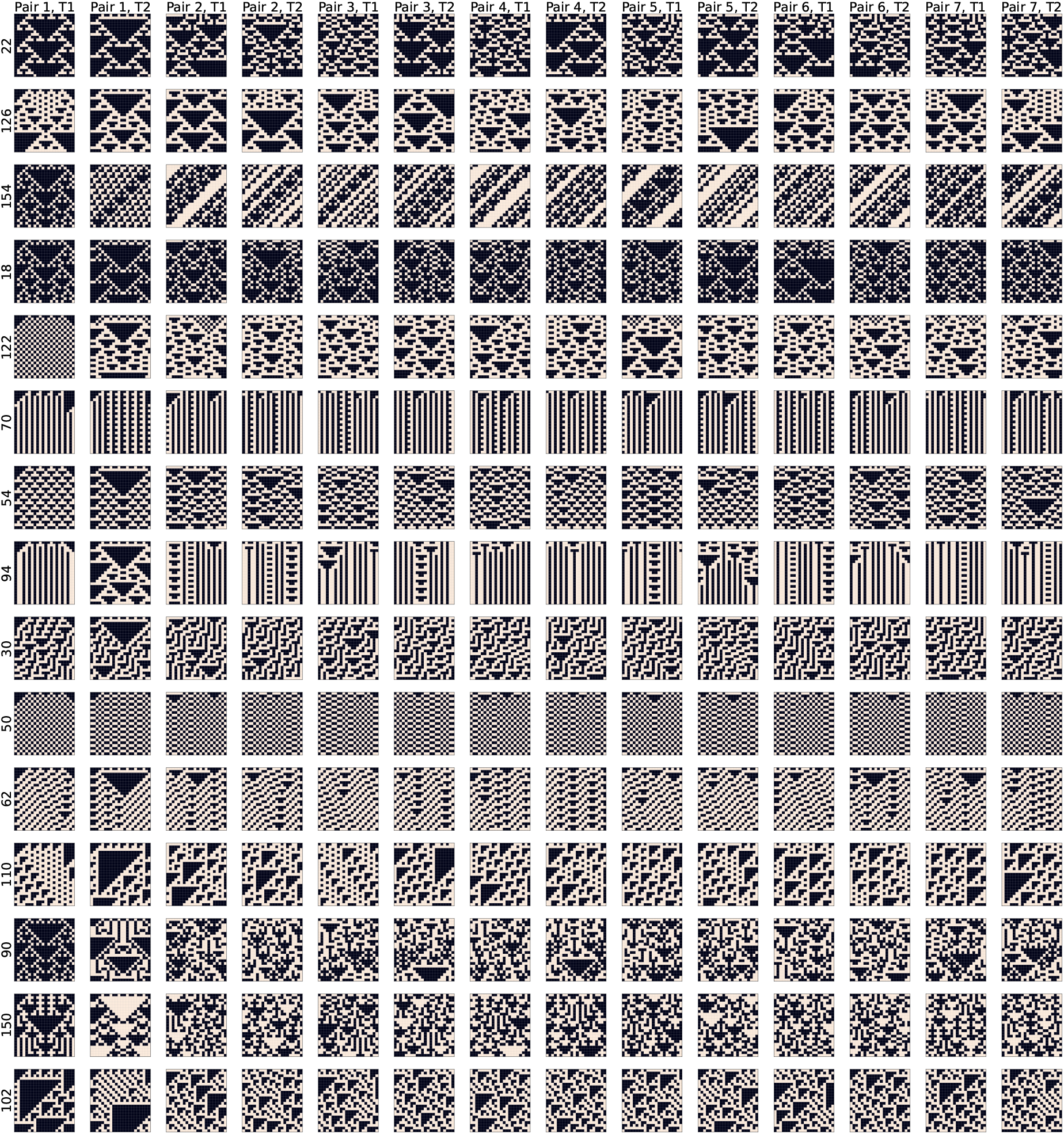
All 105 pairs of target patterns. Column: Initial condition, Row: CA rule. First two columns show initial condition pairs that were used in experiments that were analyzed in detail.

**Table 1.**
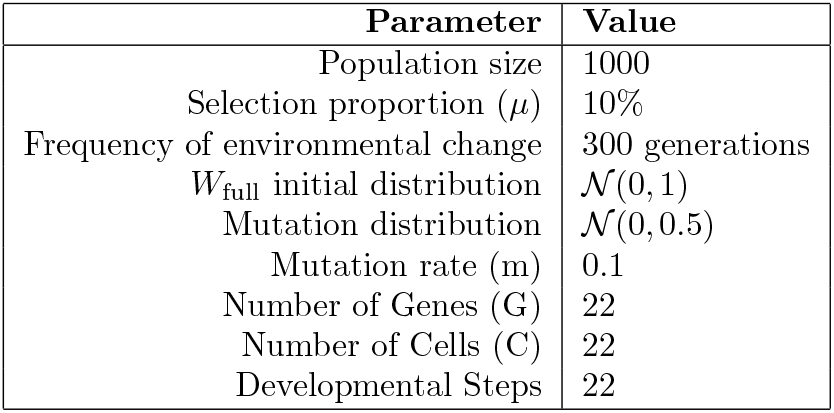
The parameters that were used in each of the experiments.

