## Supplementary material for "The variability of evolvability: properties of dynamic fitness landscapes determine how phenotypic variability evolves"

### Supplementary Information

Table 1: The parameters that were used in each of the experiments.

| Parameter | Value |
|---:|---|
| Population size | 1000 |
| Selection proportion ($\mu$) | 10% |
| Frequency of environmental change | 300 generations |
| $W_{\text{full}}$ initial distribution | $\mathcal{N}(0,1)$ |
| Mutation distribution | $\mathcal{N}(0,0.5)$ |
| Mutation rate (m) | 0.1 |
| Number of Genes (G) | 22 |
| Number of Cells (C) | 22 |
| Developmental Steps | 22 |

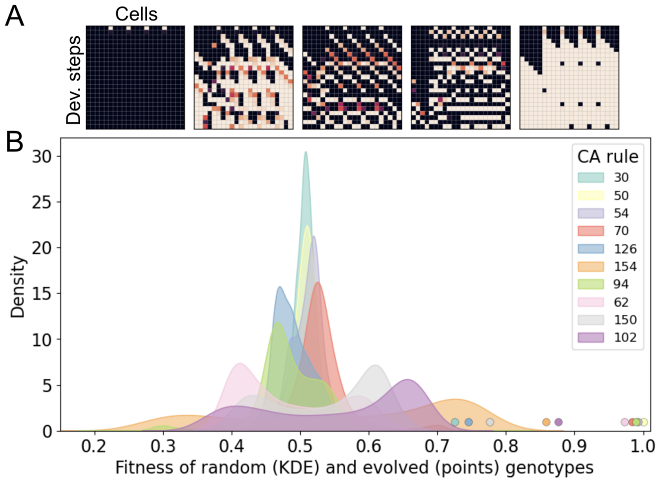


Figure 1: A) Phenotype of five randomly generated GRNs. B) Fitness distribution of randomly generated GRNs with respect to 10 different target patterns. Color coded points correspond to the maximum fitness of evolved GRNs in these fitness landscapes.

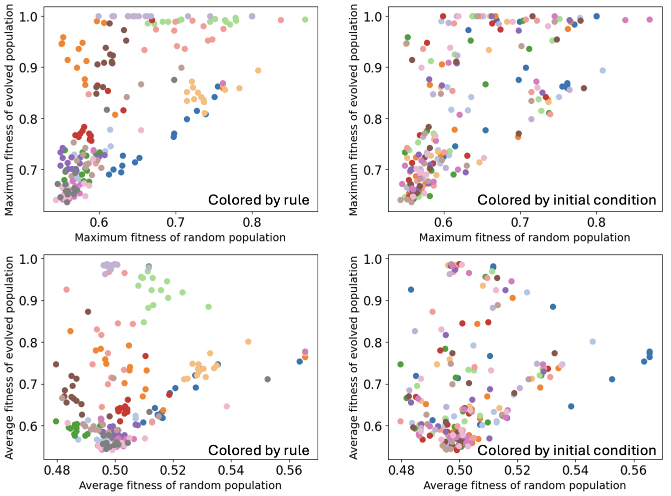


Figure 2: Maximum and average fitness of statically evolved and random populations averaged across 15 replicates for all 210 fitness landscape pairs. Left subfigures are colored by CA rule, right subfigures are colored by initial condition. Fitnesses cluster by CA rule more along both y and x axes. The standard deviation across experiments with different initial conditions is ∼3× higher than across different CA rules for evolved populations, and ∼1.5× higher for random populations, in terms of both maximum and average fitness.

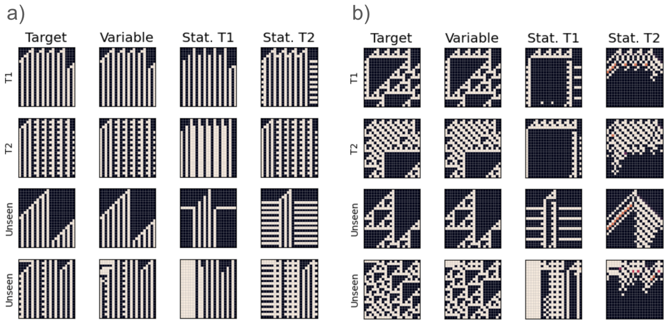


Figure 3: Example GRNs evolved to match patterns generated by rule 70 (a) or rule 102 (b). Rows: Output of the same GRN given different initial conditions. Columns left to right: Target pattern, GRN evolved in variable environment, GRN evolved in only the first environment, and GRN evolved in only the second environment. In these settings, GRNs evolved in variable environments were able to better match unseen patterns generated by the same CA rules.

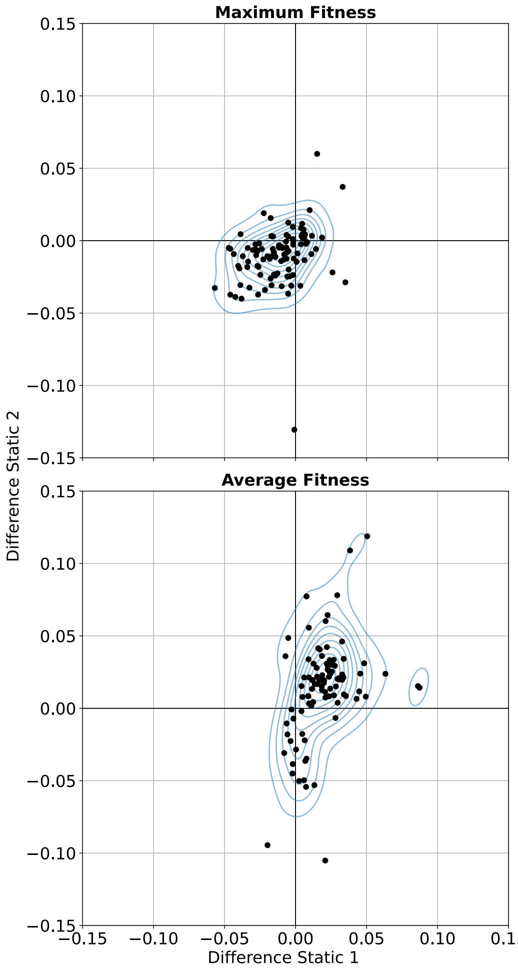


Figure 4: The effect of environmental variability on maximum and average fitness in cases where the alternative evolutionary target patterns were generated using the same initial conditions with different cellular automata rules. Thus, the alternative fitness landscapes didn't share global optima. Everything else was kept the same such that results from Fig. 2 are comparable. Again, environmental variability has a positive effect on average fitness. Additionally, environmental variability has an overall negative effect on maximum fitness. Patterns hold when accounting for multiple testing.

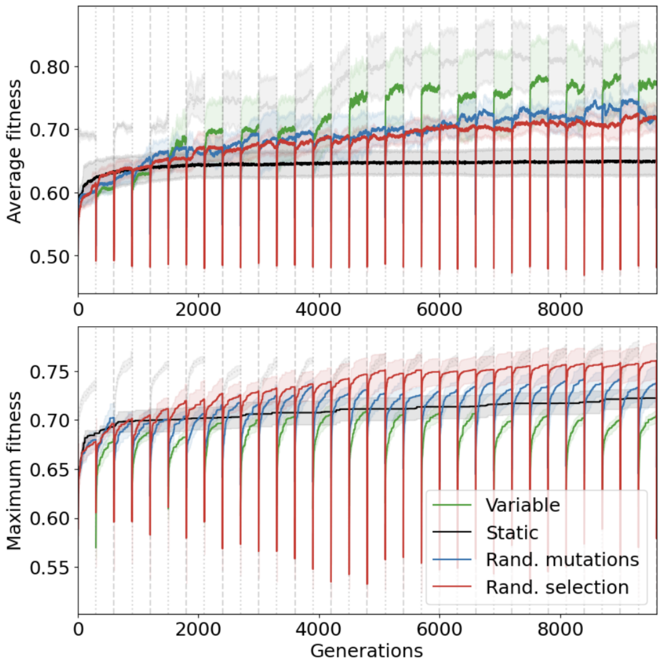


Figure 5: Average (example experiments with rule 102) and maximum (example experiments with rule 150) fitness in the population over time in variable and static environments, as well as in runs where the individuals were randomly mutated (blue lines) and in runs where the target pattern was randomized for three consecutive generations (red lines) every 300 generations. For runs with variable environments the line is grayed out for generations evolving to the alternative target for easier visual comparison. Both random mutation and random selection had a consistently positive effect over static experiments, while the effect of environmental variability was more variable.

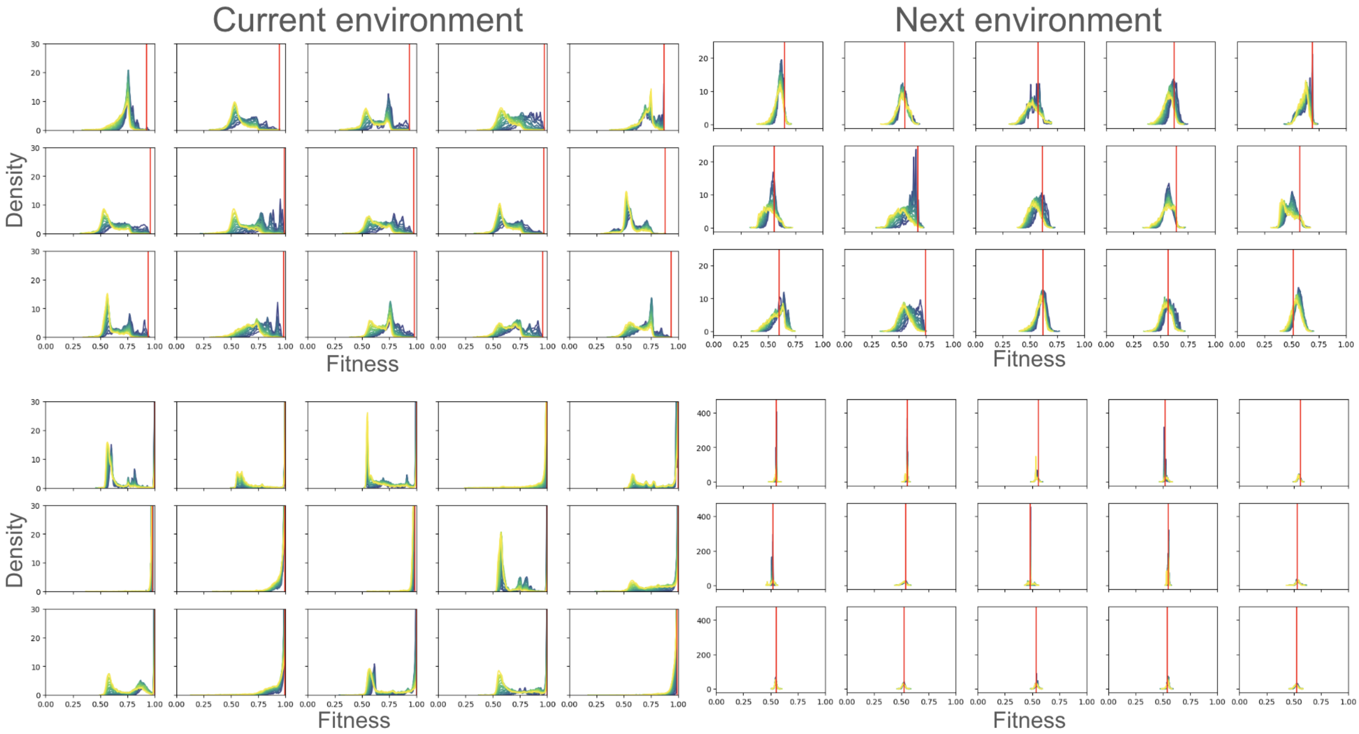


Figure 6: Distribution of fitness effects of mutations applied to the GRN with the highest fitness at the end of the first season (300 generations) across 15 replicate runs. Subfigures on the left show the fitness calculated with respect to the current, subfigures on the right with respect to the next environment. Top half shows replicate runs of experiments with rule 54, while the bottom half shows rule 122. Different colors show different mutation rates, ranging from $\mathcal{N}(0, 0.063)$ (blue) to $\mathcal{N}(0, 0.3)$ (yellow). Red lines show evolved GRNs' fitness. While most mutations had a negative effect in both experiments with respect to the current environment, in runs evolved for rule 54 (top half) a lot of these same mutations were beneficial with respect to the environment in the next generation.

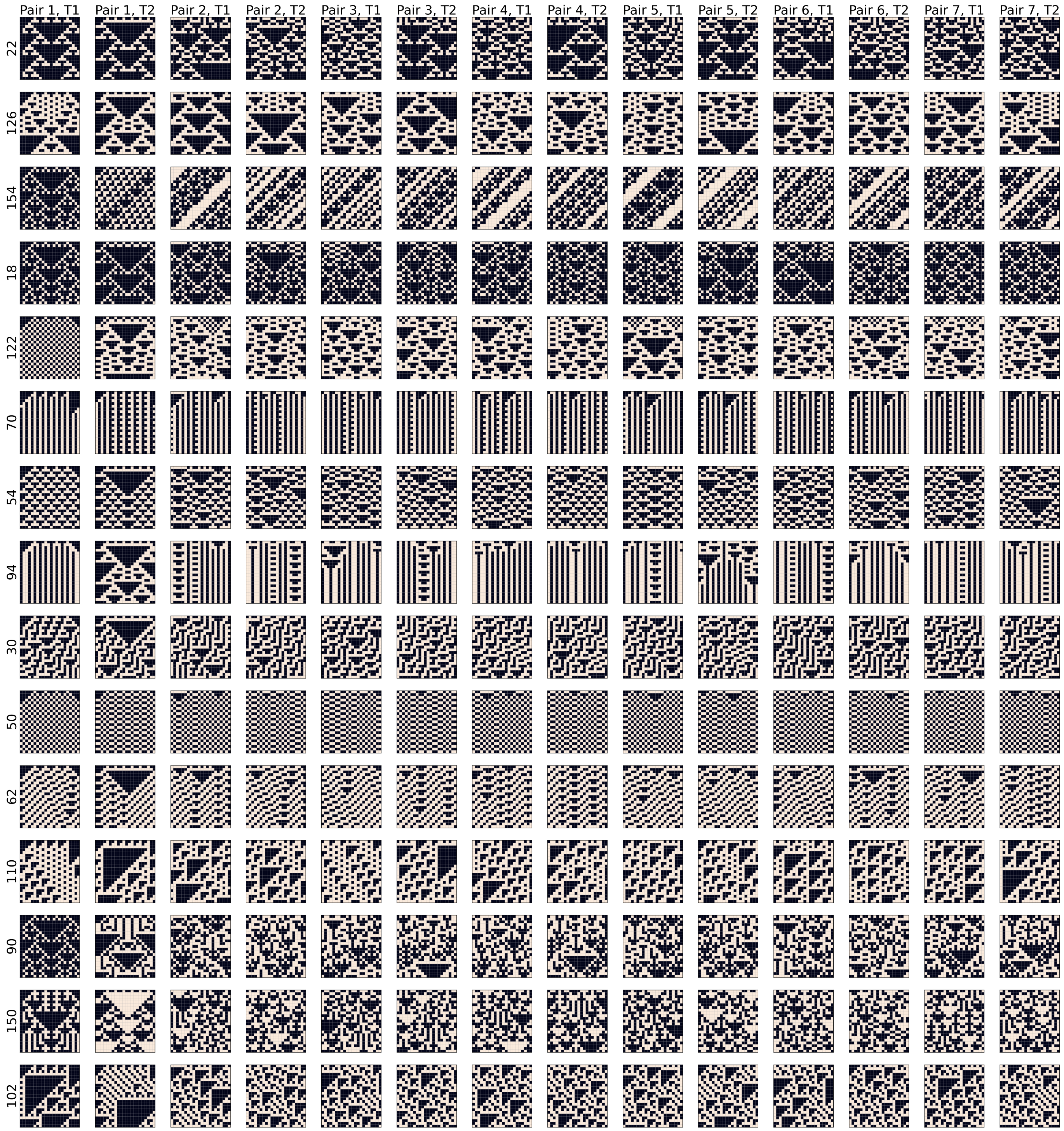


Figure 7: All 105 pairs of target patterns. Column: Initial condition, Row: CA rule. First two columns show initial condition pairs that were used in experiments that were analyzed in detail.
